# Encapsulated actomyosin patterns drive cell-like membrane shape changes

**DOI:** 10.1101/2021.10.20.465228

**Authors:** Yashar Bashirzadeh, Hossein Moghimianavval, Allen P. Liu

## Abstract

Cell shape changes from locomotion to cytokinesis are, to a large extent, driven by myosin-driven remodeling of cortical actin patterns. Passive crosslinkers such as α-actinin and fascin as well actin nucleator Arp2/3 complex largely determine the architecture and connectivity of actin network patterns; consequently, they regulate network remodeling and membrane shape changes. Membrane constriction in animal cell cytokinesis proceeds by assembly and contraction of a contractile ring pattern rich in α-actinin and myosin at the equator of the cell cortex, with which the ring is contiguous. Here we reconstitute actomyosin networks inside cell-sized lipid bilayer vesicles and show that, depending on vesicle size and concentrations of α-actinin and fascin, actomyosin networks assemble into ring and aster-like patterns. Anchoring actin to the membrane enhances the interaction of the contractile networks with lipid membrane but does not change the architecture of the patterns. A membrane-bound actomyosin ring exerts force and constricts the membrane. An Arp2/3 complex-mediated actomyosin cortex is shown to assemble a ring-like pattern at the equatorial cortex and contribute to myosin-driven clustering of the cortex and consequently membrane deformation. An active gel theory unifies a model for the observed membrane constriction and protrusion induced by the membrane-bound actomyosin networks.

## Introduction

Cell morphogenesis, shape change, and motility are highly dependent on actin networks acting in different regions of a cell ^1,2^. In the thin actin cortex under the cell membrane, the pool of actin filaments is spatiotemporally regulated by different proteins, resulting in different functionalities of the actin networks ^3,4^. The actin cortex supports the cell’s morphology and can rearrange into structures such as lamellipodia or lead to the formation of protrusions such as filopodia at the cell periphery ^5^. Actin networks at the cell periphery are organized by myosin motors, passive crosslinkers, and nucleation factors under a variety of conditions. Detachment of actin from the membrane and depletion of Arp2/3 complex both contribute to the initiation of cell protrusion at the cell periphery while short crosslinkers, such as fascin, participate in the assembly of actin bundles for cell protrusion ^6–8^. In cytokinesis, another large-scale self-organization event, myosin and α-actinin intensely localize at the cell equatorial plane to assemble the contractile ring, the positioning of which is regulated by Arp2/3 complex-mediated actin cortex ^9,10^. Simulation of actomyosin networks in confinement also showed that myosin-driven contractility has also a major role in the formation of ring-like cortex structures ^11^, which are found at the periphery of T cell immunological synapse, axons, and ring complex of podosomes ^12–14^. It was shown that actin ring-like vortices and aster structures in the cortex upon adherence of HeLa cells to a substrate are highly regulated by the Arp2/3 complex while myosin motors were not involved in these transitions ^15^.

The interplay between different actin-binding proteins in living cells makes the mechanistic study of actin network morphogenesis challenging. There has been a myriad of studies on *in vitro* reconstitution of active and passive actin networks that have unraveled the role of actin binding proteins in network self-organization. By selective activation of myosin in a pool of actin filaments on supported lipid bilayers, it has been shown that actomyosin networks contract radially towards the center of the activation region in a cooperative manner ^16^. Myosin contractility towards the barbed end of crosslinked actin filaments can induce polarity sorting and formation of aster-like structures. However, the role of myosin as an active motor or passive bundler in the formation of actin patterns depends on myosin concentration as well as crosslinker type and concentration. Myosin at low concentrations acts as an actin crosslinker in the presence of a short crosslinker fascin, while it actively contracts actin bundles to form asters at high concentrations ^17^. Myosin was recently shown to have more motility and higher velocities on fixed actin filaments bundled by fascin whereas it becomes almost immobile and trapped in a mixed polar network crosslinked by alpha-actinin ^18^. Considering the diverse set of actin binding proteins and different members of motor proteins in cells, how specific structures arise from the same pool of proteins remains unknown.

Addition of biomembranes in actin reconstitution platforms made it possible to reconstitute more sophisticated structures and study the physics and mechanism governing the interaction of actin networks with cellular membrane ^19–23^. In the absence of molecular motors, branched actin networks reconstituted on lipid bilayer membranes demonstrated the physics and mechanism governing membrane protrusion by cortical actin ^24^. In the presence of myosin motors and depending on network tension, actin cortex polymerized on the outside of giant unilamellar vesicles (GUVs) could crush the vesicle or peel the membrane ^25^. When encapsulated inside the GUVs, however, the actomyosin cortex was shown to cluster in the periphery of membrane or in the center of the vesicle depending on the strength of actin network-membrane attachment ^25^. Interestingly, this phenomenon has been investigated recently in *Xenopus* egg extract droplet in oil emulsions and is modeled as a tug-of-war between contraction waves and spontaneous bridge formation between the cluster and the membrane. It has been shown that the contraction wave period and bridge formation time scale are dependent upon the confinement size ^26^. In such a confined environment and in the presence of actin crosslinkers and crowding agents, reconstituted actomyosin bundles formed rings while increasing their contractility led to cluster formation ^27^. Contractile actomyosin rings linked to the membrane of GUVs and formed by crosslinking of actin by vinculin and talin have been reconstituted to mimic cell division ^28^. These contractile rings induce transient shape changes in GUVs and contract to a final cluster form. However, the involvement of a branched actin cortex in contractile ring positioning and membrane constriction is missing in this work despite its important role in cytokinesis. Recently, it was shown that actin clustering could occur solely due to α-actinin crosslinking activity in GUVs and that α-actinin-actin bundles could form rings or peripheral asters depending on GUV size ^23^. When reconstituted together, α-actinin and fascin sorted in separate domains to form central aster patterns in GUVs ^23^. However, the physical conditions and the contribution of individual actin crosslinkers and nucleation factors which induce the dynamic evolution of contractile actomyosin networks into diverse patterns and actomyosin interaction with the membrane in confinement have not yet been studied.

Here, force generation and confinement size-dependent pattern formation of actomyosin networks in the form of contractile rings and asters is investigated. By tuning the amount of active and passive actin crosslinkers, we determine the optimal conditions for contractile ring assembly in GUVs. By changing membrane surface properties, we then demonstrate pattern formation in the form of peripheral actomyosin rings and asters. Membrane-bound actomyosin rings exert contractile force and slightly deform GUV membrane. We then show that Arp2/3 complex-dendritic actomyosin networks in the presence of actin crosslinkers assemble contractile ring-like patterns at the equatorial plane of GUVs and condense into clusters to induce membrane deformation. An active gel model describes membrane deformation caused by both a membrane-bound actomyosin ring and cortical cluster. Our results highlight the central role of closed boundary conditions in the formation of emergent actomyosin patterns and consequently biological membrane shape changes by these patterns.

## Results

### The formation of contractile actomyosin ring and aster patterns depend on GUV size and the relative concentration of myosin and passive crosslinkers

We wanted to first establish the contractility of actomyosin networks and transmission of the contractile forces *in vitro*. To do this, we used Fourier transform traction cytometry (FTTC) ^29–31^ to measure the traction stresses exerted by actomyosin networks to a soft substrate (Young’s modulus ∼0.75 kPa) through biotinylated actin-streptavidin linkages (**Supplementary Fig. S1a, see Methods**). Non-uniform distribution of neutravidin beads bound to biotinylated actin resulted in the formation of an F-actin ‘template’ pattern upon polymerization (**Supplementary Fig. S1b-c**). Upon the addition of myosin at molar ratio of 1:20, F-actin immediately bundled, contracted, (**Supplementary video S1**), and deformed the substrate (**Supplementary videos S2-S3**), resulting in instant increase of traction stresses (**Supplementary Fig. S1d-e, Supplementary video S4**) and strain energy density to a plateau (**Supplementary Fig. S1f**). A second burst of contraction could be induced by adding more ATP (**Supplementary Fig. S1d-f**). An inverse contraction force-velocity relationship was observed in the system upon addition of myosin (**Supplementary Fig. S1g**) ^32^. Local divergence of bead displacement field upon the addition of myosin revealed a convergent contraction towards the template and actin enrichment in the template (**Supplementary Fig. S1h-j**). These results provide a quantitative means of traction stresses and work produced by actomyosin networks and show that active filaments of muscle myosin at a molar ratio of 0.05 deliver cellular-scale mechanical energy to an attached substrate at time scales on the order of seconds ^31^. The results also show that non-covalent bonds of biotin-avidin efficiently transmit actomyosin contractile forces to induce deformation.

In order to reconstitute contractile actomyosin ring, we sought to find the molar concentrations of crosslinkers and myosin that can induce formation of rings in GUVs. Knowing that our system of actin filament and myosin creates robust contraction, we first explored how actomyosin networks self-organize in GUVs in the absence of membrane attachment and how actin crosslinkers fascin or *α*-actinin influence actomyosin network organization. In the absence of actin crosslinkers, encapsulation of actomyosin networks at myosin molar ratios ranging between 0.05 to 0.0125 in DOPC/cholesterol GUVs all resulted in instant contraction and clustering of actin filaments at either the center or periphery of GUVs (**Fig. 1a, Supplementary Fig. S2**), as expected. We next encapsulated a mix of myosin, α-actinin, and fascin, each at different molar ratios, with actin inside GUVs. Irrespective of myosin concentration, addition of α-actinin did not change actomyosin condensation into a cluster (**Fig. 1b**). Myosin-fascin-actin bundles, however, formed actin meshworks with filopodia-like membrane protrusions (**Fig. 1c**), without actin condensation. In the presence of both α-actinin and fascin, when α-actinin concentration was lower than fascin concentration, stiff actin rings were formed in small GUVs (diameter < 15 µm) (**Fig. 1d, green arrows, and Fig. 1e, green arrows, Supplementary video S5**). In large GUVs (diameter > 20 µm), centripetal contraction condensed actin into clusters resulting in the formation of aster-like structures (**Fig. 1d, red arrow, and Fig. 1e, left, Supplementary video S5**) following elongation of actin bundle arms outside clusters (**Supplementary Fig. S3a-b, Supplementary video S6**). α-Actinin localized in the cluster (**Supplementary Fig. S3c-d**). Stiffer actin bundle arms outside the cluster often formed membrane protrusions (**Fig 1d, yellow arrow, and Supplementary Fig. S4**).

**Figure 1.**
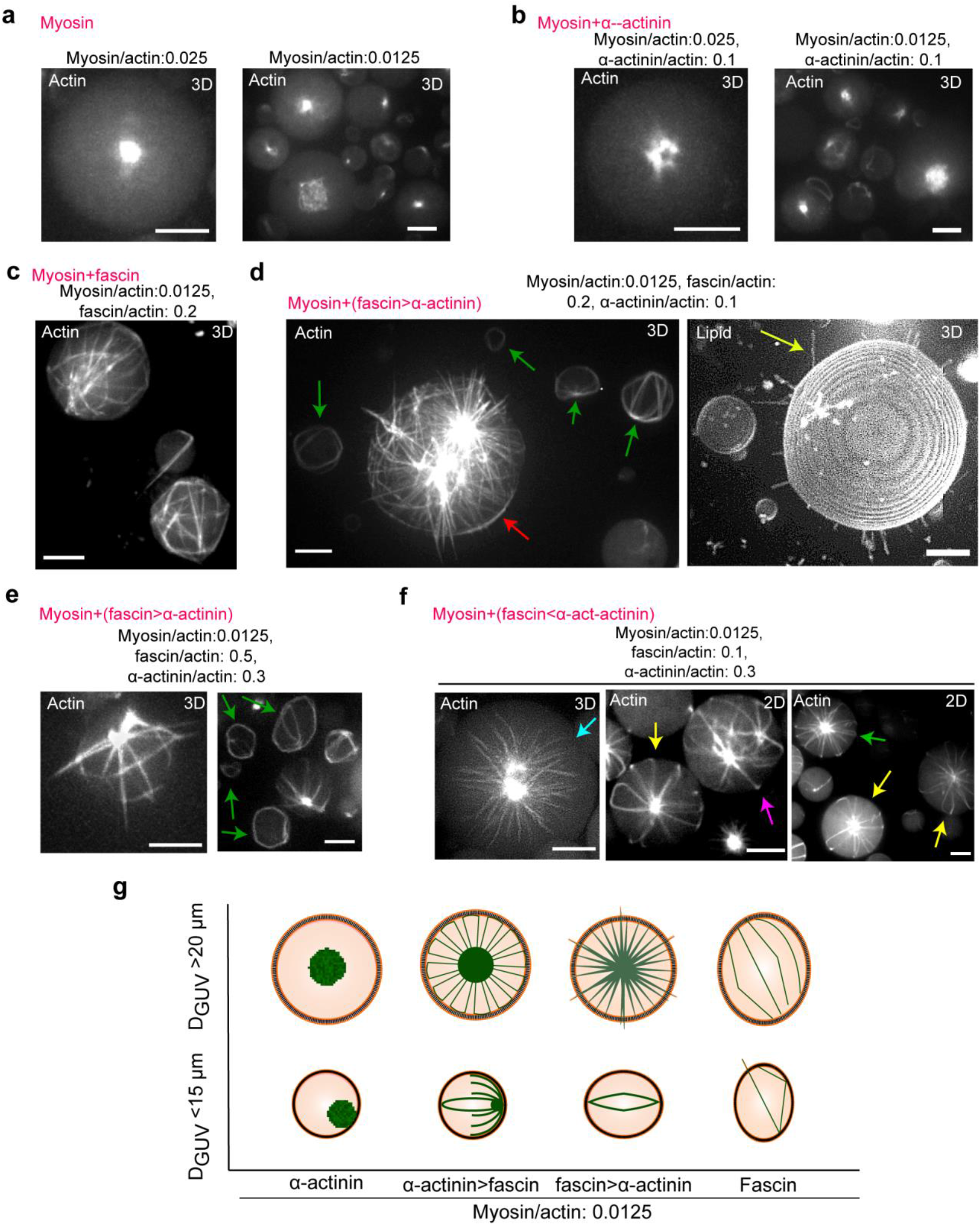
Diverse actomyosin network patterns emerge in confinement as a function of GUV size and concentration of actin crosslinkers. **(a-f)** Representative 3D reconstructed fluorescence confocal images of actin networks in the presence myosin, α-actinin, and/or fascin at molar ratios indicated. Actin, 5 μM. Scale bars, 10 μm. **(d)** Green arrows represent actin bundle rings in small GUVs. Red arrow points to a large GUV with encapsulated actin aster and ring. **(f)** Yellow arrows show actin aster with long actin bundles bent at the GUV periphery. These actin bundles turned and elongated towards GUV lumen and may appear as axially symmetric structures in 2D (green arrow). Cyan arrow shows an actin aster pattern with bundle length shorter or equal to GUV radius. Pink arrow points to two peripheral actin asters formed at two poles of a GUV. **(g)** Schematic illustration of the emergence of crosslinker type- and GUV size-dependent actomyosin network patterns.

At high α-actinin/fascin ratios, the vast majority of actin bundles formed aster-like patterns at the GUV center (**Fig. 1f**). Large actin bundles were bent at the GUV periphery and looped backed into the GUV lumen (**Fig. 1f, yellow arrows**). Almost axially symmetric structure with regular spacing between actin bundles was observed (**Fig. 1f, green arrow**). However, actin bundle arms whose radius was less than GUV radius, stayed straight (**Fig. 1f, cyan arrow**). Occasionally, peripheral actin bundle arms were joined at single or multiple clusters to form peripheral actomyosin asters (**Fig. 1f, magenta arrow**). It should be noted that, in the absence of myosin, α-actinin-fascin-actin bundles form rings and asters at high α-actinin concentrations ^23^. The majority of actin bundle rings in this condition were positioned at the equatorial plane of GUVs and parallel to the substrate a few hours post-encapsulation (**Supplementary video S7**). **Fig. 1g** summarizes the distinct actomyosin patterns that can form by tuning the concentrations of fascin and α-actinin. Together these results showed that α-actinin-actin bundles with large filament spacing contribute to actomyosin clustering, which, in turn, results in the formation of aster-like patterns in the presence of fascin. Fascin-parallel actin bundles do not cluster in the presence of low myosin concentrations, and together with a lower concentration of α-actinin, form a contractile actomyosin ring in small GUVs.

### Anchoring actin to GUV membrane localizes actomyosin patterns at the GUV periphery but does not change the architecture of actomyosin ring and aster patterns

Since contractile actomyosin rings and asters assemble in the cell cortex and interact with the plasma membrane ^9,15^, we next sought to test the formation of cortical actomyosin patterns bound to GUV membrane. Membrane binding via biotinylated actin-streptavidin/biotinylated lipid linkages resulted in the formation of peripheral patterns of actin bundles. Membrane-bound α-actinin-actin bundles formed peripheral rings (**Fig. 2a**). In the presence of α-actinin and myosin, actin filaments condensed into clusters to form peripheral aster-like patterns (**Fig. 2b**). Increasing myosin concentration further enhanced clustering at the periphery (**Supplementary Fig. S5**). Membrane-bound actomyosin asters resembled aster-like cortical actin networks of cells during cell adhesion ^15^. Replacing α-actinin with fascin induced the formation of peripheral actomyosin meshworks (**Supplementary Fig. S6, Supplementary video S8**), in stark contrast to unanchored fascin-actomyosin system (**Fig. 1c)**. Complete suppression of fascin-induced membrane protrusions is again in line with recent observations that membrane attachment reduces actin network protrusions in cells ^7^. The observed fascin-actomyosin meshworks were also reported to form in bulk at low myosin and high fascin concentrations ^17^. Like unanchored actomyosin networks, our observations indicate that fascin facilitates the formation of actomyosin bundles while α-actinin contribute to myosin-driven cluster formation in membrane-bound actomyosin networks. Like unanchored networks, reconstitution of membrane-bound actomyosin networks in the presence of both fascin and α-actinin resulted in the formation of stiff aster-like patterns. However, cluster was located at the GUV periphery and packed bundles elongated into the GUV lumen with suppressed membrane protrusion (**Fig. 2c**). In agreement with this result, enhanced actin-membrane interaction in the cell was recently shown to reduce plasma membrane protrusion ^7^. The phenotype of aster-like patterns in the absence (**Fig. 2d**) and presence (**Fig. 2e**) of fascin is schematically illustrated.

**Figure 2.**
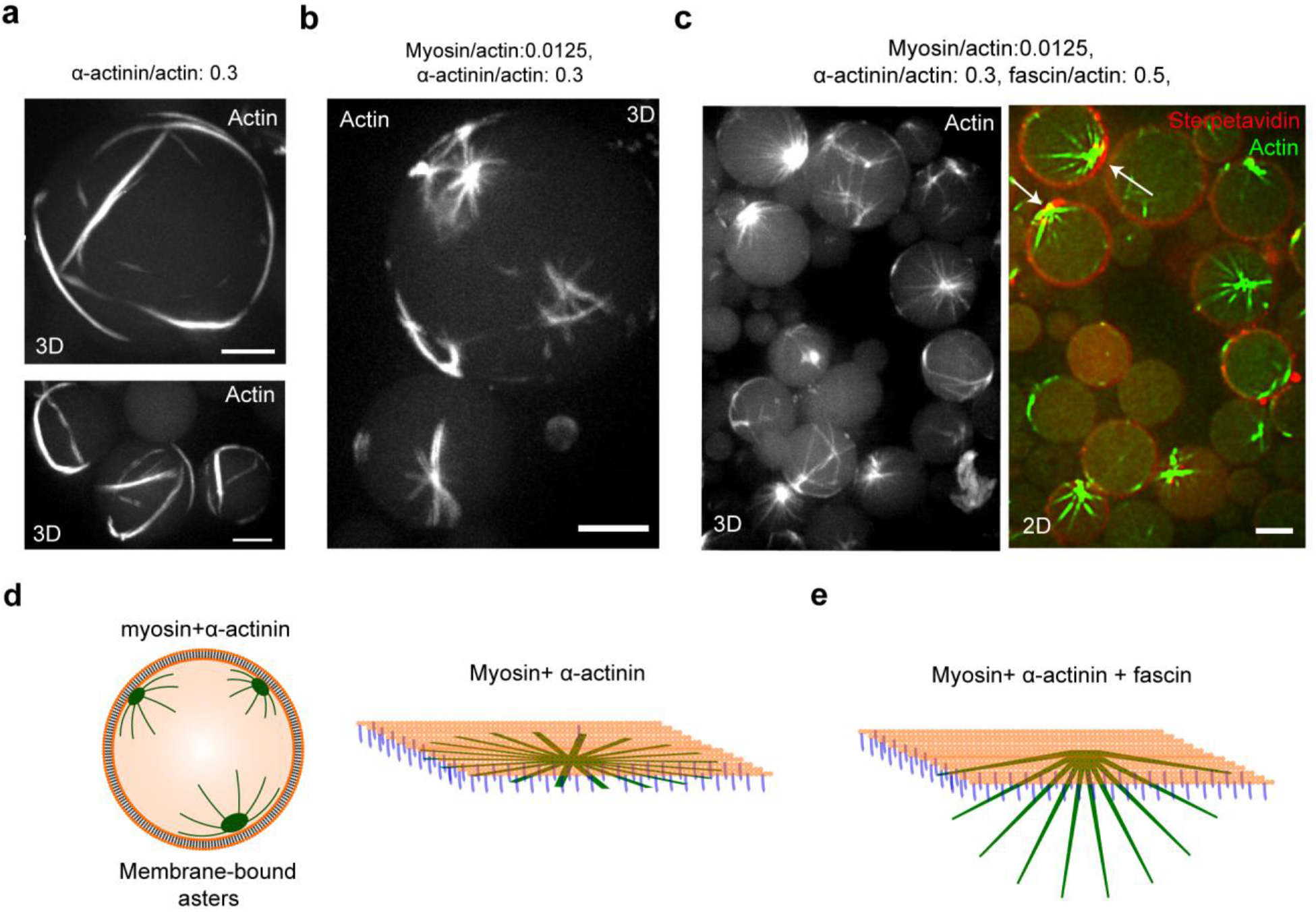
Enhanced actin-membrane interaction does not change the architecture of contractile actin patterns but suppresses actin bundle protrusion. **(a)** Representative 3D reconstructed fluorescence confocal images of membrane-bound actin bundles in the presence of α-actinin at the molar ratios indicated. Actin, 5 μM. Scale bars, 10 μm. **(b)** Representative 3D reconstructed fluorescence confocal images of peripheral actin asters formed in the presence of myosin and α-actinin at molar ratios indicated. Actin, 5 μM. Scale bars, 10 μm. **(c)** Representative 3D reconstructed fluorescence confocal images of membrane-bound actin in the presence of myosin, α-actinin, and fascin at molar ratios indicated. White arrows show aggregated actin-streptavidin linkages at the site of actin cluster. Actin, 5 μM. Scale bars, 10 μm. **(d)** Schematic illustration of clustering of membrane-bound actin networks and the emergence of short actin asters in the presence of α-actinin and myosin. **(e)** Schematic illustration of the effect of fascin on the architecture of cortical actomyosin patterns in large GUVs (diameter > 20 µm). While attachment to GUV membrane suppresses protrusion, α-actinin and fascin together form aster structures elongating into the lumen of GUVs.

### Membrane-bound actomyosin ring patterns deform GUV membrane during self-assembly

Since our long-term goal is to generate a contractile ring in a GUV to mimic cytokinesis, we wanted to understand better how these actomyosin rings form. Using a high fascin molar ratio in the presence of α-actinin, a condition we found to form rings in small GUVs, we successfully captured time evolution of the assembly of a membrane-bound contractile actomyosin ring over ∼ 40 minutes. Similar to the formation of contractile bundles on soft substrates, we first noticed the formation of a thin actin bundle template in the form of a ring initially (**Fig. 3a, Supplementary video S9**). During contractile ring assembly we noticed that GUV membrane was constricted (**Fig. 3a, red arrows, and Supplementary video S10, dashed box**) at the locus of accumulated actin in the ring. By measuring membrane angles with respect to the plane of actomyosin ring (**Fig. 3b**), and assuming membrane tensions, σ_1_ and σ_2_ are equal to their average, σ ^33^, we found the maximum contractile force of the ring (F_c_), measured at time point 16 min to be 5.89 × 10 ^-6^ σ [N]. Membrane tension for DOPC GUVs of similar size (5 µm in radius) is estimated to be ∼ 3× 10 ^-5^ [N/m] under iso-osmotic conditions ^34^. Using this value, contractile force of the ring would then be ∼176 [pN], which is comparable to 390 [pN] generated by sliding contractile ring in fission yeast protoplasts ^33^. Concurrent to force generation, actin localization increased over time to complete ring assembly (**Fig. 3c-d**). By analyzing the fluorescence intensity of α-actinin and myosin along several line regions of interest that intersect the actomyosin ring, we found that α-actinin and myosin were not entirely co-localized and can partially segregate along the ring (**Fig. 3e-f)**, similar to aster-like patterns (**Supplementary Fig. S3c**).

**Figure 3.**
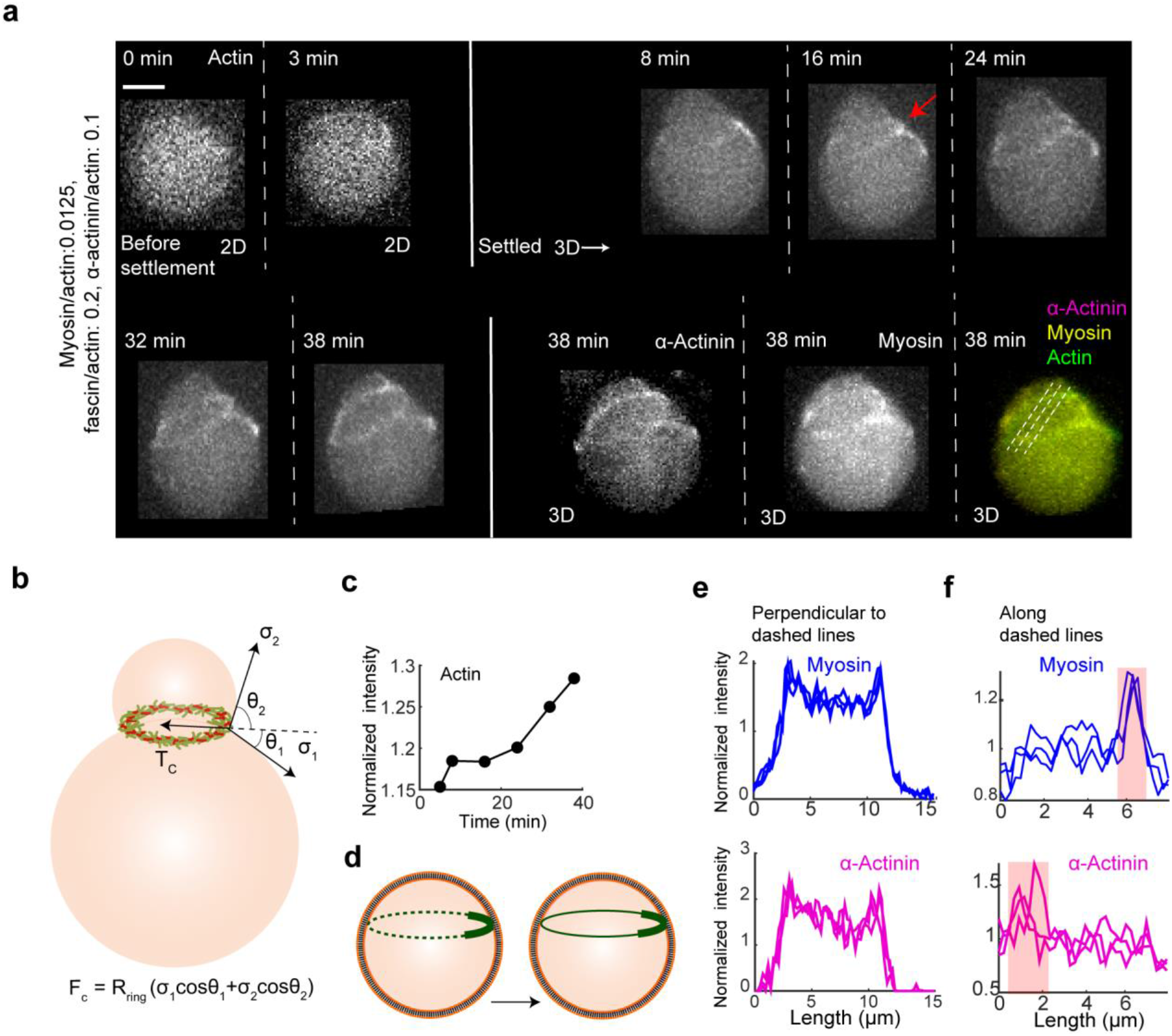
Membrane-bound contractile ring patterns deform GUV membrane. **(a)** Representative fluorescence confocal actin images showing a time-lapse of contractile ring assembly in the presence of myosin, fascin, and α-actinin as well as 3D confocal images of α-actinin, myosin, and merged images of the ring at the last time point (38 min). At the first two time points, actin images were captured as single 2D images before GUV settlement. The red arrow shows the region where maximum ring constriction occurs. Dashed lines show the line along which actin, myosin, and α-actinin intensities were quantified. Actin 5 μM. Myosin/actin, 0.0125 (M/M). α-Actinin/actin, 0.3 (M/M). Fascin/actin, 0.5 (M/M). Scale bars, 10 μm. **(b)** Schematic illustration of parameters used to calculate contractile force (F_c_) as a function of membrane tensions (σ_1,_ σ_2_). **(c)** Average values of maximum actin intensity along the 3 white dashed lines in (a). Intensity values were normalized to average intensity along each line. **(d)** Schematic illustration of myosin-induced clustering and contractile ring assembly in a GUV. **(e-f)** Intensity profiles of α-actinin and myosin along perpendicular bisector of **(e)** and along **(f)** the 3 white dashed lines in (a) at the last time point (38 min). Intensity values were normalized to average intensity along the line.

### Dendritic actomyosin cortex in the presence of actin crosslinkers assemble into a ring-like contractile pattern at the GUV equator and induce membrane deformation

Actomyosin ring contraction and membrane constriction in the cell is highly regulated by a branched actomyosin cortex under the membrane and adjacent to the ring. To test the role of branched actin cortex on the organization of actomyosin patterns and contractile ring assembly, we reconstituted an Arp2/3-branched dendritic actin cortex on the inner surface of GUVs in the presence and absence of myosin, α-actinin, and fascin. Arp2/3 complex was activated by membrane-bound constitutively active VCA domain of neural Wiskott Aldrich syndrome protein. The dendritic actin patterns formed an actin shell at the GUV periphery regardless of the presence of α-actinin and fascin (**Supplementary Fig. S7**). Inclusion of myosin, and together with the concentrations of α-actinin and fascin known to form rings (**Fig. 1g**), in the dendritic networks caused symmetry breaking and actin clustering at the periphery and the emergence of small spherical membrane protrusions (**Fig. 4a-c**). A myosin molar ratio of 0.0125 was sufficient for symmetry breaking, and consequently clustering actin towards the membrane due to strong attachment of actin to the membrane ^25,35^. Interestingly, however, actin was intensely localized at the equatorial plane of GUVs in the form of a ring-like pattern (**Fig. 4c**). 3D intensity analysis of the contractile cortex structures showed that the contractile cortex not only promote the formation of ring-like patterns at the equatorial cortex (**Fig. 4d-g**) but tend to induce actin clustering, often in multiple directions (**Fig. 4f, red arrows**). To rule out the existence of artifacts in the formation of ring-like patterns, we also analyzed 3D intensities of GUV membrane and dendritic actin cortex structures formed in the absence of mysoin and actin crosslinkers and confirmed that actomyosin cortex structures indeed induce the formation of the observed ring-like patterns (**Supplementary Fig. S8**). Membrane deformation was frequently observed at the location of actin clusters, which often appeared in the form of a ring-like pattern at the neck of the deformed membrane (**Fig. 4d, white arrow**). The mode and time-scale of membrane deformation resembled the constriction of membrane-bound actomyosin bundles formed in the presence of talin and vinculin ^28^. We occasionally observed large membrane protrusions at the site of actomyosin clusters in the active networks (**Fig. 4h**). The absence of actin cortex in these membrane protrusions suggested the possibility of bleb formation due to the generation of cytoplasmic pressure induced by cortex tension in the reconstituted active networks ^36^. Together these data demonstrated that myosin-driven contractility contributes to both the formation of ring-like patterns at the GUV equator and a build-up of cortex tension for membrane protrusion in confinement.

**Figure 4.**
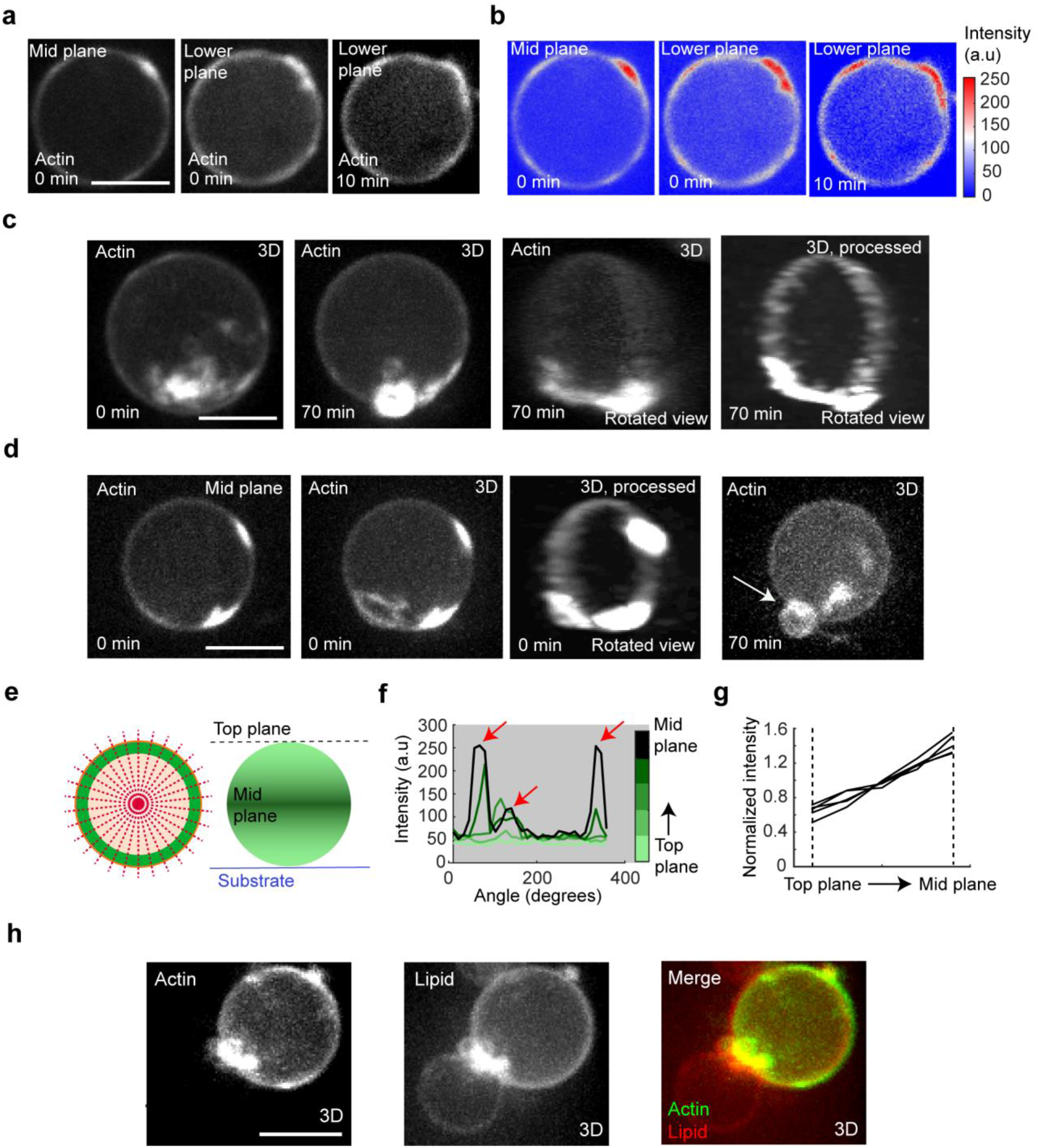
Dendritic actomyosin networks in the presence of actin crosslinkers self-assemble into a ring-like equatorial cortex. **(a)** Representative fluorescence confocal actin images of a crosslinked actomyosin cortex 0 min and 10 min after imaging. Representative 3D reconstructed fluorescence confocal images of a membrane-bound actomyosin cortex in the presence of VCA, Arp2/3 complex, α-actinin, and fascin at different time points. **(b)** Color map of actin intensities in **(a). (c)** Representative 3D reconstructed fluorescence confocal actin images of a crosslinked actomyosin cortex 0 min and 70 min after imaging. **(d)** Another example of a 3D reconstructed fluorescence confocal actin images of a crosslinked actomyosin cortex 0 min and 70 min after imaging. White arrow points at the ring-like cluster around the neck of the deformed membrane. **(e)** Schematic representation of 2D top view (left) and 3D side view (right) of an encapsulated actin cortex with highly enriched actin at its mid plane. The red dashed lines represent radial lines along which maximum actin intensities are plotted in **(f). (f)** Actin cortex intensity around equally spaced circular *z*-planes from the top plane to mid plane of the GUV in **(d)**. Actin intensity (*y*-axis) at each plane (color map) was measured by detecting maximum intensity along 30 radial lines (red dashed lines in **(e)**) drawn with an equal angular interval (*x*-axis). Red arrows point at the intensity of actin clusters at the GUV equatorial plane. **(g)** The mean value of normalized actin cortex intensity from top plane to mid plane of 6 GUVs. Normalized mean intensity (*y*-axis) for each cortex was measured by taking the mean value of maximum intensities along 30 radial lines (red dashed lines in **(e)**) from top plane to mid plane of each GUV (*x*-axis) with *z*-interval of 2.4 µm averaged over the mean value at each *z*-plane. **(h)** Representative fluorescence confocal images of actin, lipid, and merged lipid-actin for a GUV with a bleb-like protrusion. Actin: 5 µM, Arp2/3 complex: 1 µM; His_6_-tagged VCA: 0.5 µM, α-actinin: 0.5 µM, and fascin: 1 µM for all panels. Scale bars, 10 μm.

## Discussion

We encapsulated membrane-bound and –unbound actin networks in the presence of myosin motors and passive crosslinkers, α-actinin and fascin, and showed that diverse actomyosin patterns in the form of clusters, rings, and asters emerged, the architecture of which depend on confinement size and the concentration of the motor and crosslinkers. Although α-actinin forms flexible actin bundles in the form of rings in confinement, it promotes entrapment of myosin and clustering of actin bundles. α-actinin-actomyosin clustering and disassembly to monomers could thus facilitate the elongation of parallel fascin-bundles and consequently the formation of aster patterns ^37^. Parallel fascin-actin bundles are less prone to motor-induced clustering which, at certain concentrations of α-actinin and myosin, gain enough flexibility to promote contractile ring assembly. Membrane-bound contractile ring could deform the lipid membrane during self-assembly, but it lacks a branched actomyosin cortex to position, stabilize, and build tension in the membrane. Arp2/3 complex-mediated branched actin networks in the presence of fascin have been shown to form clusters with short actin bundle arms in the form of aster-like structures but protrusion of bundles are inhibited when the networks are attached to GUV membrane in the form of a dendritic actin cortex ^38–40^. Similarly, here, a dendritic actomyosin cortex formed an actin shell at the periphery. In the presence of α-actinin and fascin, the dendritic actomyosin cortex intensely localized at the equatorial plane in a ring-like pattern. The simulation of α-actinin-actomyosin networks in confinement showed that high actin treadmilling rate shifts myosin-driven centripetal actin clustering towards the periphery which consequently results in the formation of a ring-like actomyosin cortex. It was further shown, in simulations and cultured T cells that inhibition of treadmilling induces collapse of ring-patterns into clusters. Here, actomyosin clustering occurred more frequently in the plane of ring-like actomyosin patterns in the cortex. Peripheral collapse of the actomyosin networks into clusters rather than centripetal clustering was indicative of strong attachment of the cortex to the membrane. Actomyosin collapse and disruption of the cortex are expected to cause cytoplasmic pressure buildup and consequently bleb-like protrusion of the membrane as observed at the site of cluster ^41–43^. The presence of a ring-like actin pattern at the neck of the deformed bilayer also suggests the presence of a constricting force which may contribute to bleb formation and equilibration. Thus, as there was no evidence of membrane detachment at the site of the cluster, it is expected that actomyosin networks contribute to membrane deformation in two ways, by cortex tension and generation of excess cytoplasmic pressure, and by exertion of force on the membrane.

An elastic model of bleb formation has successfully described the effect of actomyosin cortex tension on Laplace pressure and consequently bleb expansion ^44^. To account for membrane constriction at the site of protrusion, here, we include a force, F_c_, acting in the plane of constriction and opposing GUV total tension (cortex plus membrane tension), σ_g_, and protruded membrane tension, σ_b_ (**Fig. 5**). Given the balance of forces in the cortex and fluid pressure balance between the GUV and bleb at equilibrium, one would obtain σ_g_ = 1.41 σ_b_ + c, where c is the term corresponding to the elastic resistance of the cortex to its contraction and is negligible at equilibrium in the absence of cytoplasmic elasticity ^44^. Since GUV cytosol is mostly made up of water, and lacks crowded intracellular structures as opposed to a cell, it lacks elastic resistance thereby causing the term elastic resistance to vanish (i.e. c→0), which corresponds to the convergence of GUV cortex tension towards equilibrium ^44^. Balance of forces at the plane of constriction results in F_c_ = 2.75 × 10^−6^ (0.917 σ_g_ + 0.819 σ_b_) [N]. Strikingly, at σ_g_ = 1.44 σ_b_ (compare with σ_g_ = 1.41 σ_b_ + c), F_c_ equals the maximum contractile force (F_c_ = 5.88 σ_b_ × 10^−6^) generated by the actomyosin ring at the same myosin concentration (**Fig. 3**), and also holds the assumption of negligible cortex elastic resistance at equilibrium, i.e. c ∼ 1.44 σ_b_ - 1.41 σ_b_ = 0.03 σ_b_. The ratio (σ_g_ / σ_b_ = 1.44) between membrane tension of branched actomyosin-bound lipids (σ_g_) and bare lipids (σ_b_) is very close to the ratio of tension (∼1.3) between a GUV doublet covered with a branched actomyosin cortex and its initial tension before myosin addition and continuation of actin polymerization ^45^, although, the cortex tension at the site of constriction can differ from the tension of floppy membrane domains caused by the presence of a corral domain and Marangoni effect ^46^. The presence of dendritic actin cortex formed by Arp2/3 complex is the major difference between constriction force at the site of the bleb and the force generated by actomyosin ring in cortex-less GUVs. Because myosin drives constriction, and crosslinking forces have negligible effect on the contractile force ^33^, our results suggest that Arp2/3 complex participates in bleb formation, yet has minimal effect on constriction at the bleb site. Our findings together show that inherent physical properties of actin binding proteins in confinement are sufficient to assemble minimal, yet cell-like, actomyosin patterns that drive membrane shape changes.

**Figure 5.**
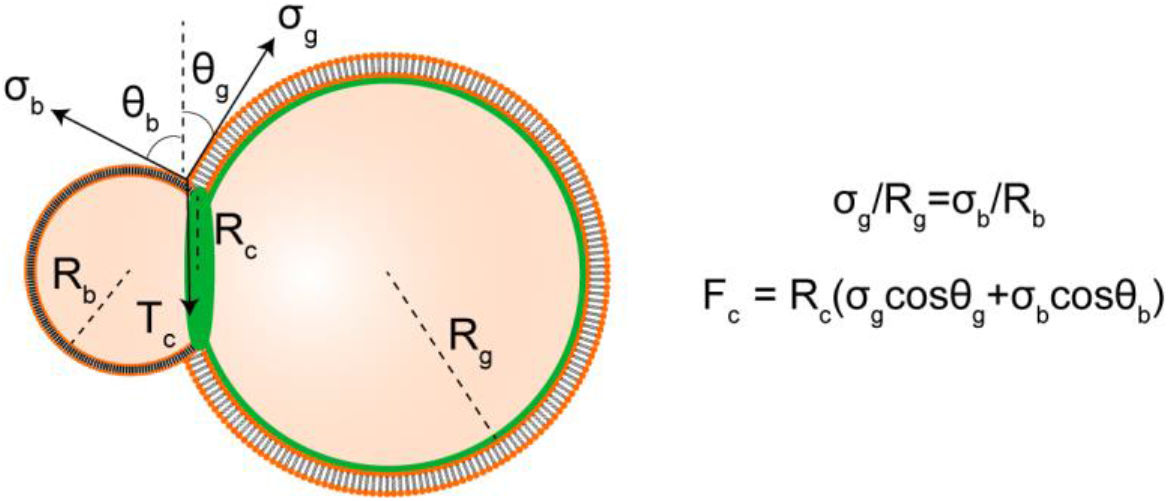
A physical model explains how cortex tension and clustering induce GUV deformation. The schematic illustrates parameters used for modeling membrane deformation and bleb formation induced by actin cortex contraction and clustering.

## Methods

### Proteins and Reagents

Actin, α-actinin, and Arp2/3 complex were purchased from Cytoskeleton Inc, USA. ATTO 488 actin was purchased from Hypermol Inc, Germany. Fascin was purified from *E. coli* as Glutathione-S-Transferase (GST) fusion protein ^23^. TMR-α-actinin was gifted by David Kovar (University of Chicago). Myosin was labeled using Alexa Fluor™ 647 microscale protein labeling kit (Thermo Fisher Scientific). We purified hexa-histidine-tagged VCA (His_6_-tagged VCA) domain from neural Wiskott Aldrich syndrome protein following the same steps described previously ^38^.

### Measurement of Contractile Stresses via Traction Force Microscopy

Traction stresses exerted by actomyosin networks were quantified by using Fourier transform traction cytometry (FTTC) ^29–31^. Actin filaments were coupled to a soft Matrigel (Young’s modulus ∼ 0.75 kPa and Poisson’s ratio ∼ 0.5) allowing the measurement of contractile stresses exerted as traction forces on the gel. For this purpose, 200 nm yellow-green neutravidin-coated beads (Invitrogen) were embedded on the top surface of a soft gel following previous protocols ^47,48^. The substrate was prepared in an open chamber, allowing the addition of actin and other components (i.e., myosin and extra ATP) for time-lapse measurements. Monomeric actin (including 10% Alexa Fluor 568 actin) and 10 mol % biotinylated actin in polymerization buffer was added to the chamber. Monomeric actin (including 10% Alexa Fluor 568 actin) with 10 mol % biotinylated actin in polymerization buffer were added to the chamber. Coupling of biotinylated actin to the fluorescent neutravidin beads enabled the transmission of contraction forces to the substrate. The displacement field of the embedded fluorescent beads enabled the extraction of exerted traction forces using FTTC. Strain energy was measured using custom-written MATLAB routines ^49^.

### GUV Generation

Lipids containing 70% 1,2-dioleoyl-sn-glycero-3-phosphocholine (DOPC), and 30% cholesterol in a 1:4 mixture of mineral oil and silicone oil was first made in a glass tube. The total concentration of lipids was 0.4 mM in the oil mixture. For membrane-bound actomyosin networks (without VCA and Arp2/3 complex), lipid mixture was composed of 69% DOPC, 30% cholesterol, and 1% 1,2-dioleoyl-sn-glycero-3-phosphoethanolamine-N-(cap biotinyl) (sodium salt) (biotinyl cap PE). For dendritic actomyosin networks (with VCA and Arp2/3 complex), lipid mixture was composed of 70% DOPC, 25% cholesterol, and 5% 1,2-dioleoyl-sn-glycero-3-[(N-(5-amino-1-carboxypentyl)iminodiacetic acid)succinyl] (nickel salt) (DGS-NTA(Ni)). For labeling GUVs, 0.1% of 1,2-dioleoyl-sn-glycero-3-phosphoethanolamine-N-(lissamine rhodamine B sulfonyl) (Rhod-PE) was added to the lipid mixture. All lipids were purchased form Avanti Polar Lipids. Mineral oil and silicone oil were purchased from Sigma-Aldrich.

Then, 5 μM actin, including 10% ATTO 488 actin and 4% biotinylated actin (in the case of membrane-bound networks without VCA and Arp2/3 complex), in polymerization buffer (2 mM MgCl_2_, 4.2 mM ATP, 0.2 mM CaCl_2_, and 50 mM KCl, in 15 mM Tris, pH 7.5) and 7.5% OptiPrep was prepared and kept in ice for 15 min. Myosin (0.06-0.25 μM), α-actinin (0.5-1.5 μM), fascin (0.5-1.5 μM), VCA (1 μM), Arp2/3 complex (1 μM), or their combinations was then added to the sample. Our previous observations found that, in contrast to reconstituted dendritic networks outside GUVs, high molar ratios of Arp2/3 complex with respect to actin (∼0.2) were needed for the formation of cortical dendritic patterns inside GUVs ^38^.

GUVs were produced by a modification of the cDICE method ^23,50^. A rotor chamber was 3D-printed with clear resin and mounted on the motor of a stir plate and rotated at around 1,200 rpm. 0.7 mL outer solution (200 mM glucose matching the osmolarity of inner solution) and 5 mL of lipid-in-oil dispersion were sequentially transferred into the rotating chamber. The difference in density between the two solutions results in the formation of two distinct layers with a water/oil interface. Then, a water-in-oil emulsion was created by adding 0.7 mL of lipid/oil mixture to 20 μL of protein mixture and rigorously pipetting up and down 6-7 times. The emulsion was then pipetted into the rotating chamber. It should be noted that the prepared mixture of actin binding proteins was added to the actin solution 2-3 seconds before encapsulation in droplets, and droplets traveled through the lipid dispersion in the rotating chamber. As the droplets cross the water-oil interface in the rotating chamber, they form a bilayer and are released in the outer solution as GUVs.

### Microscopy and image analysis

GUVs were transferred to a 96 well plate for microscopy. Images were taken using an oil immersion Plan-Apochromat 60 x/1.4 NA objective on an inverted microscope (Olympus IX-81) equipped with an iXON3 EMCCD camera (Andor Technology), AOTF-controlled lasers (Andor Technology), and a Yokogawa CSU-X1 spinning disk confocal. Acquisition of images was controlled by MetaMorph (Molecular Devices). Single and z-stack images of lipid and actin were captured with 561 nm excitation at exposure time of 20-25 ms and 488 nm excitation at exposure time of 350-500 ms, respectively. 3D images were reconstructed by brightest point projection of z-stack image sequences in Fiji/ImageJ. 3D processed images in **Fig. 4c-d** were generated using Fiji/ImageJ complemented with the plugin Squassh ^51^. Intensities in **Fig. 4f** were obtained using the Fiji/ImageJ plugin, ‘Oval Profile’. Using this plugin, maximum intensities (cortical actin intensity in actin images and membrane intensity in lipid images) were detected along 30 radii from the center to the points of a circle surrounding the GUV of interest from the top plane to mid plane (equatorial plane) of the GUV (**Fig. 4e**).

## Acknowledgements

We thank David Kovar (University of Chicago) for TMR-α-actinin. This work is supported by the National Science Foundation (1844132 and 099332) and National Institutes of Health (NIBIB EB030031-01) to A.P.L..

