## Supplemental information for "Encapsulated actomyosin patterns drive cell-like membrane shape changes"

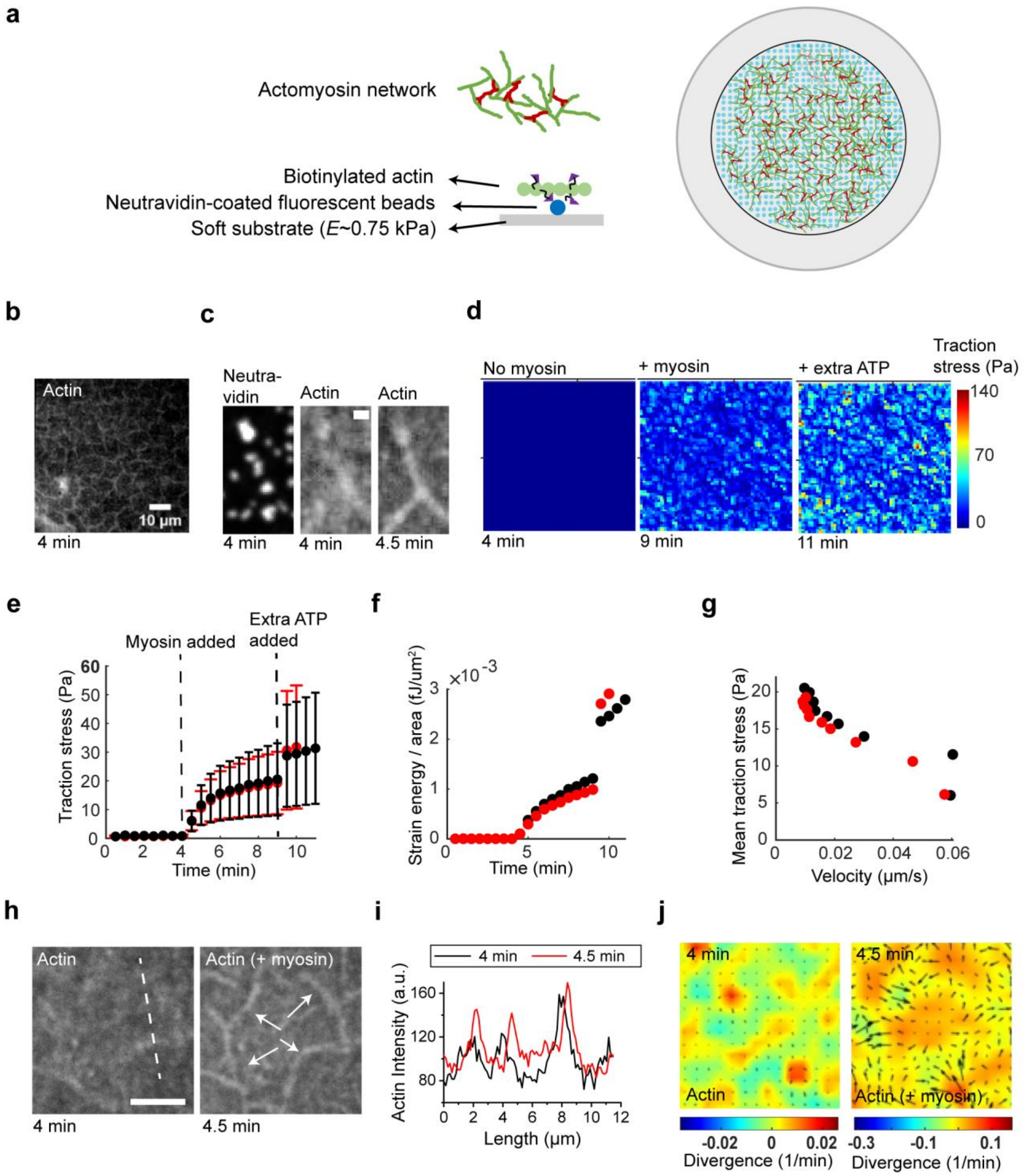


**Figure S1.** Contractility and exertion of traction forces by actomyosin networks. (a) Schematic illustration of reconstitution of actomyosin networks on a soft gel. 3 μM actin in polymerization buffer containing 1.2 mM ATP was polymerized and anchored to neutravidin-coated fluorescent beads embedded in the top surface of a soft substrate. (b) A representative field of view from fluorescence confocal images of reconstituted actin networks. (c) The traction stress field determined from bead displacements at 3 different times after the start of imaging: 4 min (left, right before adding 0.15 μM myosin), 9 min (right before adding additional 2.2 mM ATP), and 11 min (the final stage). (d) Representative confocal fluorescence images of neutravidin (left) and actin (middle and right) in a small region at the indicated time points. The images show that F-actin have more presence in discrete regions where neutravidin-coated beads are localized. (e and f) Time evolution of mean traction stress (e) and strain energy over area (f) for two substrates (black and red) with the same stiffness. Black represents the field shown in (b). Error bars indicate standard deviation. (g) Traction stress-bead velocity relationship in the contractile networks. (h) Neutravidin beads under traction stress are sheared towards local areas rich in F-actin. Representative fluorescence confocal images of actin right before and after addition of myosin in an area of interest is shown on the left. (i) Actin intensity profiles along the white dashed before (black) and after (red) addition of myosin are shown. (j) Divergence of bead displacement field in the region at the two time points (before and after addition of myosin) is shown. Vector field in divergence maps shows bead displacement.

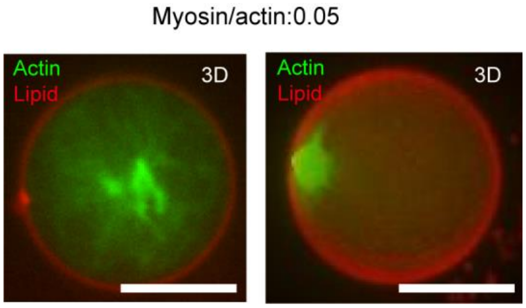


**Figure S2.** Representative 3D reconstructed fluorescence confocal images of actin (green) clustered by myosin at the center (left) or periphery (right) of GUVs (red).

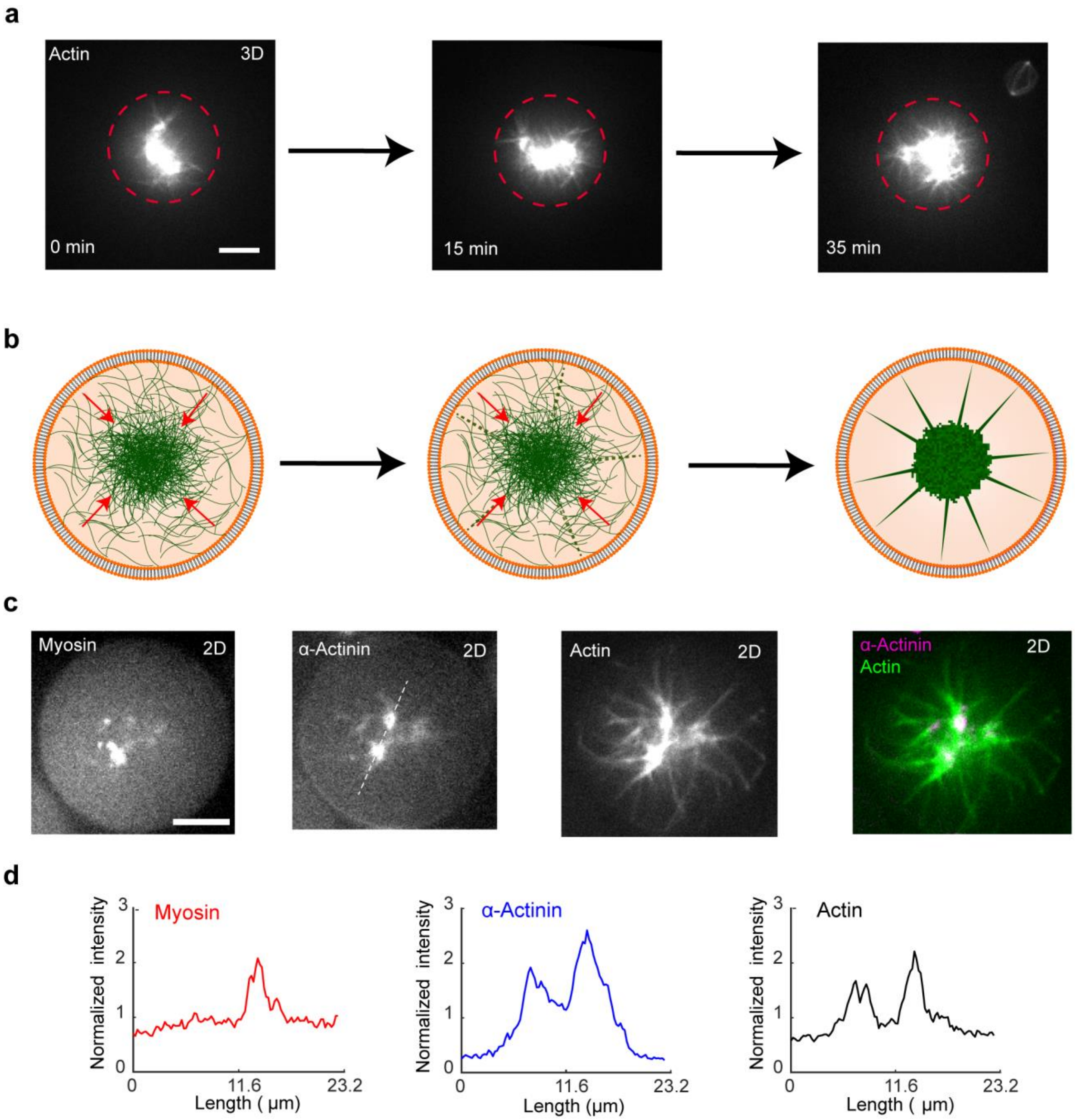


**Figure S3.** The dynamics of the formation of aster-like actomyosin patterns. (a) Representative 3D reconstructed fluorescence confocal actin images of an encapsulated actomyosin network at different time points. Actin, 5 μM. Myosin/actin, 0.0125 (M/M). α-actinin/actin, 0.3 (M/M). Fascin/actin, 0.5 (M/M). Scale bars, 10 μm. (b) Schematic illustration of the assembly of aster-like actin patterns in confinement. (c) Representative confocal fluorescence images of myosin, α-actinin, actin, and merged α-actinin and actin in an encapsulated aster-like pattern. Actin, 5.8 μM. Myosin/actin, 0.0125 (M/M). α-actinin/actin, 0.3 (M/M). Fascin/actin, 0.5 (M/M). Scale bars, 10 μm. (d) Intensity profiles of myosin, α-actinin, and actin along the dashed line in (c). Intensity values were normalized to average intensity along the line.

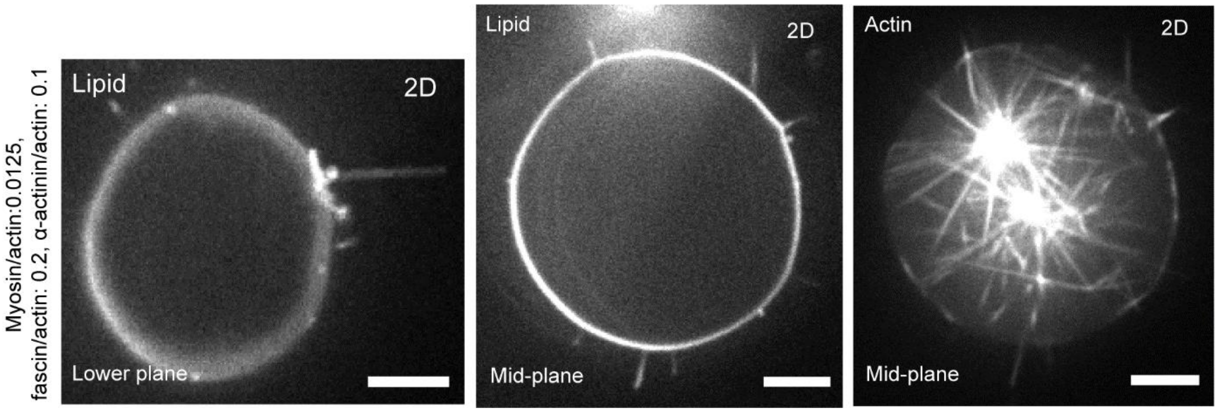


**Figure S4.** 2D fluorescence confocal images of aster-like actin pattern, shown in Figure 1d, at three different *z*-planes.

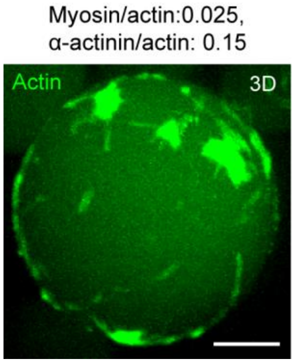


**Figure S5.** A representative 3D reconstructed fluorescence confocal image of peripheral actin asters formed in the presence of myosin and α-actinin at molar ratios indicated. Actin, 5 μM. Scale bars, 10 μm.

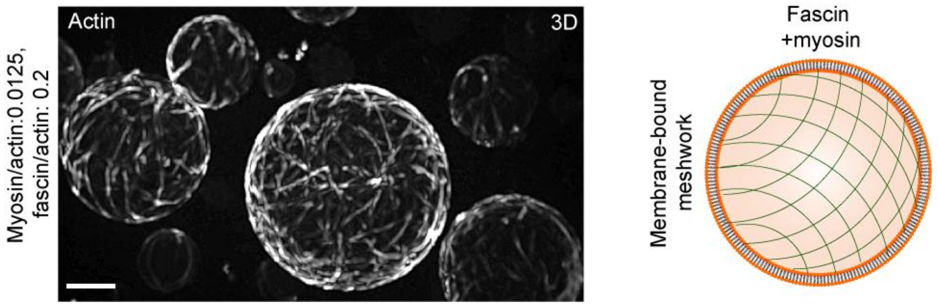


**Figure S6.** Representative 3D reconstructed fluorescence confocal images (left) and schematic illustration of membrane-bound actin meshworks (right) formed in the presence of myosin and fascin at molar ratios indicated. Fascin enhances the formation of actomyosin rings and meshwork, by inhibiting myosin-driven clustering. Actin, 5 μM. Scale bars, 10 μm.

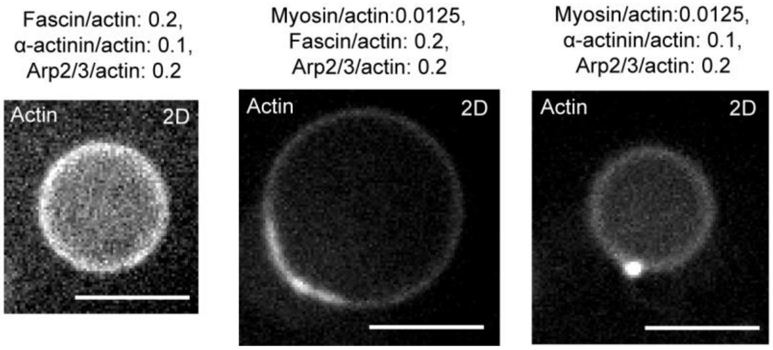


**Figure S7.** Representative 2D fluorescence confocal images of dendritic actin cortex in GUVs in the presence of actin binding proteins at molar ratios indicated.

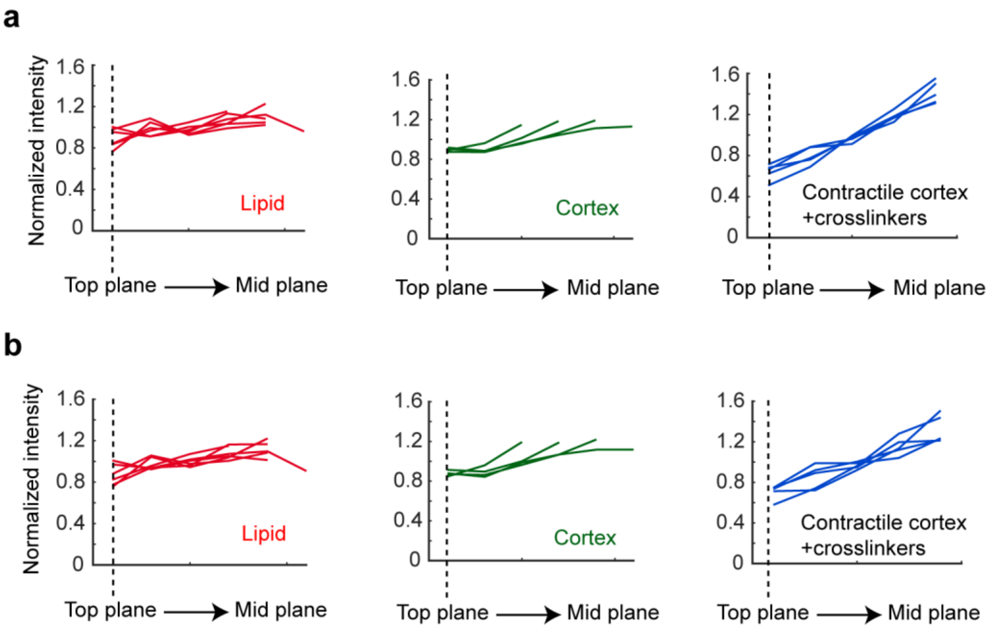


**Figure S8.** The average (a) and median (b) of normalized maximum intensity along radial lines shown in Fig. 4e drawn from top *z*-plane to mid *z*-plane of 5 GUVs (red), actin images of Arp2/3 complex-cortex in 4 GUVs (green), and contractile cortex in the presence of α-actinin and fascin in 6 GUVs (blue). Each individual line represents a different GUV.

**Supplementary videos:**

**Video S1.** Time-lapse image sequence of the contraction of actin networks shown in supplementary Figure 1b.

**Video S2.** Time-lapse image sequence of neutravidin-coated fluorescent beads embedded to the substrate of actin networks shown in Video S1.

**Video S3.** Time-lapse image sequence of the neutravidin-coated fluorescent beads in Video S2 including displacement vector field of the beads.

**Video S4.** Time-lapse images of the magnitude of traction stresses exerted by contractile actomyosin networks (also see Supplementary Figure 1d).

**Video S5.** z-stack image sequence taken from myosin-α-actinin-fascin-actin bundle networks shown in Figure 1d (red arrow).

**Video S6.** Time-lapse actin image sequence from the formation of the central actomyosin aster shown in Supplementary Figure S4a.

**Video S7.** Rotated view of a representative 3D reconstructed fluorescence confocal image of a large aster-like structure formed in the presence of α-actinin, and fascin in a sealed chamber between two cover slips. α-Actinin / actin, 0.3. Fascin / actin, 0.1. Actin, 5 μM. Scale bar, 10 μm.

**Video S8.** Rotated view of a representative 3D reconstructed fluorescence confocal image of encapsulated actin rings formed in the presence of α-actinin, and fascin. α-Actinin / actin, 0.3. Fascin / actin, 0.1. Actin, 5 μM. Scale bar, 10 μm.

**Video S9.** Z-stack image sequence taken from fascin-bundled actomyosin meshworks shown in Figure 5.

**Video S10.** Time-lapse actin image sequence from the formation of the contractile ring illustrated in Figure 2.

**Video S11.** Actin images at time points 8 min and 16 min illustrated in Figure 2a. Maximum membrane constriction occurred in the area inside the dashed box. Alignment of time-lapse 3D images distorted image frames but membrane constriction is apparent within the dashed box.
